# Telomeric lncRNA TERRA localizes to stress granules in human ALT cells

**DOI:** 10.1101/2024.06.18.599513

**Authors:** Luca Larini, Elena Goretti, Eleonora Zulian, Emma Busarello, Stefano Maria Marino, Mona Hajikazemi, Katrin Paeschke, Toma Tebaldi, Emilio Cusanelli, Katarina Jurikova

## Abstract

TERRA, the lncRNA derived from the ends of chromosomes, has a number of well-described nuclear roles including telomere maintenance and homeostasis. A growing body of evidence now points at its role in human cells outside of nucleus—it has been found to be a component of extracellular vesicles, a player in inflammation signalling and its capacity for translation has been shown. In this work, using a combination of sensitive microscopy methods, cellular fractionation, proteomics and transcriptome analysis, we demonstrate directly for the first time that TERRA is present in the cytoplasm of human telomerase-negative cells, especially upon various stress stimuli, and that it associates with stress granules. Confirming the presence of TERRA in the cytoplasm, our work fills an important gap in the field, and contributes to the discussion about the role of TERRA as a transcript involved in nucleo-cytoplasmic stress communication.

## Introduction

The localization of long non-coding RNAs (lncRNAs) is often tightly linked to their function. In different cellular compartments, lncRNAs may exhibit distinct roles that are regulated by different environmental conditions, including their interaction partners ^1–3^.

Upon transcription in the nucleus, many lncRNAs are translocated to the cytoplasm to fulfill various roles. Nuclear export of RNAs is mediated by a dedicated export machinery, and the particular pathway may depend on the RNA characteristics. The export of lncRNAs—RNAs that are long, with one or few exons, A/U-rich and m6A-modified—is preferentially mediated by the nuclear RNA export factor 1 (NXF1) ^4^. Many lncRNAs exhibit dual nuclear and cytoplasmic localization or are shuttled to the cytoplasm upon environmental change or stress and play an important role in cytoplasmic processes such as signaling, translation regulation and stress response ^2^.

Approximately 40 % of cellular lncRNAs were found associated with polysomes, which may be linked to their degradation ^5^ rather than to productive translation ^6^. Another destination for cytoplasmic lncRNAs are the membrane-less, phase-separated organelles, such as stress granules (SGs). SGs form in response to stress-induced translation inhibition and contain a number of RNA and RNA-binding proteins (RBPs) that may vary in different conditions, including lncRNAs ^7, 8^. While the interplay between lncRNA recruitment and the formation of the cytoplasmic phase-separated bodies has not been broadly elucidated, the RNA-RNA interactions have been proposed to contribute to the formation and stabilization of these structures ^9, 10^.

One of the lncRNAs with canonically nuclear roles is the telomeric repeat-containing RNA TERRA ^11–14^. TERRA transcription is initiated from promoters in the subtelomeric region, running towards the end of the chromosome, resulting in a transcript with length spanning from 200 nt to 10 kb that carries a 5’ subtelomeric sequence as well as a telomeric sequence in its 3’ end. Vast majority of chromosome ends give rise to TERRA molecules ^15^ and they have been shown to be post-transcriptionally modified: most molecules carry the 5’ 7mG cap, ∼7% has been estimated to be polyadenylated ^16^ and recent works have also shown the presence of m6A modification ^17, 18^. The presence of the telomeric UUAGGG repeats enables TERRA to form an RNA G-quadruplex secondary structure *in vitro*, although its relevance in vivo remains debated ^19^.

TERRA has a number of well-described roles in telomere biology, including the maintenance of telomeric heterochromatin and telomerase regulation ^12, 20–25^. In addition, TERRA plays an important role in the telomerase-independent, alternative lenghtening of telomeres (ALT): its levels are upregulated in ALT cancer cell lines ^26–30^, and the R-loops it forms by pairing with telomeric DNA stimulate the recombination-based telomere maintenance typical for ALT cells ^28, 31, 32^. However, an increasing number of reports hints at its relevance outside of the nucleus. Live-cell imaging and RNA-FISH revealed TERRA re-localization to the cytoplasm in budding yeast during diauxic shift, a phase in the yeast culture growth when glucose is depleted from the media ^33^. Furthermore, TERRA has been identified as a component of inflammation-inducing exosomes in a number of murine and human cell lines ^34, 35^. A recent work has demonstrated its role in MAVS-dependent inflammation triggering in senescing fibroblasts, where TERRA and mitochondria-associated Z-DNA binding protein (ZBP1) were shown to be indispensable for the signalling activation ^36^. The transfection of TERRA-mimicking oligonucleotides in human cells leads to its translation into dipeptide repeat proteins; and antibodies raised against such proteins can be used to visualize a signal in ALT cells, opening a possibility of TERRA being translated in this system ^37^. To date, a direct observation of TERRA in the cytoplasm of human cells, elucidation of its nuclear export and further insight in its possible cytoplasmic role are missing.

In this work, we set out to study cytoplasmic TERRA transcripts in cancer cells. Using single molecule RNA-FISH, cell fractionation and dCas13-based visualization, we demonstrate for the first time that TERRA can be detected in the cytoplasm of human ALT cell lines. Furthermore, we show that the levels of cytoplasmic TERRA, similarly to nuclear TERRA, are increased in the presence of telomere DNA damage. Supported by the proteomic interactome screen and RNAi-based down-regulation, we provide evidence that NXF1 is involved in the nuclear export of TERRA. We further show that a subpopulation of cytoplasmic TERRA co-localizes with stress granules during sorbitol and sodium arsenite-induced stress. Our results indicate that TERRA, an lncRNA with well-studied nuclear roles linked to telomere maintenance, may serve alternative, cytoplasmic roles in cells with telomerase-independent, ALT telomere maintenance in response to telomere damage and during stress.

## Materials and methods

### Cell culture, transfection, transduction and stress induction

AGS (stomach adenocarcinoma cell line) were a kind gift from Christian Baron (Université de Montréal, Canada). WI-38 VA-13 (ALT human lung, SV40-transformed cell line) were obtained from ATCC. SA-OS2 (ALT human osteosarcoma cell line) was obtained from Cytion (formerly CLS; Germany). SA-OS2 cells were maintained in DMEM/F12 with 10 % FBS (Gibco), 2 mM L-glutamine (Gibco) and 50 U/ml penicillin-streptomycin (Gibco), the other cell lines in DMEM (Gibco) supplemented as above, and incubated at 37°C with 5% CO_2_. Cells were regularly tested for mycoplasma contamination.

The plasmids used for the dCas13-based visualization of TERRA were dPspCas13b-Suntag-svFC-sfGFP-eDHFR-3xFlag, pC0043-PspCas13b-sgTERRA-22nt ^38^, kindly gifted by Huaiying Zhang and Meng Xu, Carnegie Mellon University. For the depletion of POT1, we used PSuperPuroRetro-1660(hPOT1) (a gift from Christopher Counter, Addgene #11135)^39^. For the NXF1 depletion, we replaced the POT1-targeting sequence in PSuperPuroRetro-1660(hPOT1) digested with PsiI and HindIII with a NXF1-targeting hybridized insert (shNXF1pSuper-FW, 5’-CCGGCCAGTTCTGAAGAGATCCAAACTCGAGTTTGGATCTCTTCAGAACTGGTTTTTG-3’ and shNXF1pSuper-REV (5’-AATTCAAAAACCAGTTCTGAAGAGATCCAAACTCGAGTTTGGATCTCTTCAGAACTGG-3’).

PSuperPuroRetro-1660s(hPOT1)^39^ (a gift from Christopher Counter, Addgene #11136) was used as a scramble control in both experiments.

Plasmids were transfected in cells with JetPrime (Polyplus) transfection reagent or oligofectamine (ThermoFisher Scientific) in 6-well plates per manufacturer’s protocol and for all shRNA plasmids, the cells were grown in DMEM with 0.75 µg/ml puromycin.

Plasmids pLPC-TRF2-WT and pLPC-TRF2-deltaB-deltaM ^40^, a gift by Titia de Lange (Addgene #18002, Addgene #18008), were used to produce lentiviral vectors in HEK 293T cells, and the vectors were transduced in U2OS cells using 8 µg/µl polybrene.

To induce stress granule formation, the U2OS cells were incubated with 0.4 M sorbitol in supplemented DMEM for 4 hrs at 37°C, 5% CO_2_ or, alternatively, with 100 µM sodium arsenite in DMEM for 1 hr at 37°C, 5% CO_2_ and fixed immediately upon treatment.

### Single molecule inexpensive FISH (smiFISH)

The protocol was performed as described by ^41^, with details specified in ^42^. Enzymatic treatment was performed on fixed cells after permeabilization, with 200 µg/µl Purelink RNase A (Thermo Fisher Scientific) and 100 U/µl RNase T1 at 37 °C for 2 hours. For imaging, we used Nikon AX laser scanning confocal microscope with Plan Apo λ 60x Oil objective and Nikon Eclipse Ti2 microscope equipped with the CREST Optics X-Light V2 Spinning Disk module with 100X/1.4 NA oil objective.

### SmiFISH combined with immunofluorescence

For immunofluorescence, in addition to the smiFISH protocol, after the post-hybridization washes and before DAPI staining, cells were blocked for 1.5 hours with 2% BSA in PBS with 0.1% Triton-X (PBT) and subsequently incubated for 1.5 hours with primary antibodies (1:250 mouse anti-G3BP1 (sc-365338, Santa Cruz Biotechnology); 1:250 mouse anti-FUS (sc-47711, Santa Cruz Biotechnology); 1:500 mouse anti-p-histone H2A (Ser139) (05-636, Merck); 1:500 rabbit anti-TERF2IP (anti-RAP1) (NB100-292, Novus Biologicals)) washed 5 times with PBT, incubated for 1.5 hours with secondary antibodies (1:1500 goat anti-mouse conjugated with Alexa488, A28175 or 1:1500 goat anti-rabbit conjugated with Alexa647, A21245), and again washed 5 times with PBT.

The image analysis was performed in ImageJ with an in-house macro ^25, 42^ using the DiAna co-localization analysis plug-in. The shuffle function of DiAna plug-in was used for the assessment of non-randomness of G3BP1-TERRA co-localization events. Data visualization and statistics were performed in DATAtab (https://datatab.net/) or the Gnumeric software.

### dCas13-sgTERRA imaging

Plasmids were transfected in the U2OS cells as described above, in total amount of 2 µg of DNA per 35 mm well, in 1:1 dCas13:sgRNA w/w ratio. dCas13-GFP-transfected cells (without sgTERRA) were used as a negative control. Cells were fixed for imaging 48 hours post-transfection. Only dCas13-positive foci co-localizing with smiFISH TERRA signal were considered specific.

### Cell fractionation

U2OS, SA-OS2 and WI-38 VA13 cells were fractionated following the REAP protocol ^43^, using the REAP buffer with 0.1 % NP-40 for U2OS and SA-OS2 and 0.08% NP-40 for WI-38 VA13. The fractions were collected from four 100 mm plates with 80–90% confluency in ∼300 µl REAP buffer. The fractionation efficiency was verified with Western blot, using cytoplasm-specific (ɑ-tubulin) and nucleus-specific (fibrillarin) markers.

### RNA extraction & Northern blotting

Total RNA and fractional RNA was extracted with TRIzol (Thermo Fisher Scientific) per manufacturer’s instructions. For Northern blotting, 5 µg of RNA per sample were treated by incubation with 1 µl DNase I (ThermoScientific) for 1 hr at 37°C, denatured by incubation @65°C for 5 minutes, and separated in a 1.2 % agarose-MOPS gel with 1.9% formaldehyde. 3 µg of RNA treated with 2 µl PureLink RNase A (ThermoScientific) for 1 hr at 37°C were used as a control. The capillary blotting was performed overnight, onto an Amersham Hybond N+ hybridization membrane (Cytiva) in 20x SSC (0.15 M sodium citrate, 1.5 M sodium chloride). The wet membrane was cross-linked at 120 mJ/cm^2^ using a CL-1000 Ultraviolet Crosslinker (UVP). It was subsequently pre-hybridized in Church buffer (1mM EDTA, 1% (w/v) BSA, 0.5 M sodium phosphate monobasic, 7% (w/v) SDS, pH 7.2) at 42°C for at least 1 hour and hybridized with ^32^P-labeled (CCCUAA)_5_ probe at 42°C overnight. The membrane was then washed with 0.2% (w/v) SDS and 2x SSC (2 times) and 0.2% (w/v) SDS and 0.2x SSC (4 times), exposed to phosphorus screen and acquired with Typhoon laser scanner (Cytiva).

### Protein extraction & Western blotting

For isolation of protein from cells in culture, cells were collected by scraping and centrifugation for 3 minutes at 1500 g, washed with ice-cold PBS and the pellet was resuspended in 50 µl PBS. For both collected cells and aliquots from fractionation, 50 µl of the suspension was mixed with 10 µl of 6x RIPA buffer (final 1x concentration: 150 mM NaCl, 1% NP-40, 0.5% sodium deoxycholate, 0.1% sodium dodecyl sulphate, 50 mM Tris pH 7.4, 1 mM DTT, 1 mM PMSF), triturated, incubated for 20 min on ice and centrifuged for 15 min at 15000g at 4°C. The protein concentration in the supernatants was determined by Bradford assay. For Western blotting, we used primary antibodies anti-ɑ-tubulin (1:1000, sc-8035, Santa Cruz Biotechnology), anti-fibrillarin (1:800, #2639, Cell Signalling), anti-NXF1 (1:150, sc-136095, Santa Cruz Biotechnology) and anti-actinin (1:1000, sc-17829, Santa Cruz Biotechnology). As secondary antibodies, we used goat anti-rabbit IgG HRP (1:3000, ab6721, Abcam) and rabbit anti-mouse IgG HRP (1:3000, ab6728, Abcam). The signal intensity was quantified by using the Volume Tool in Image Lab 6.1 software (Bio-Rad).

### SG transcriptome analysis

Three different studies on stress granules (with different stress-inducing agens, sodium arsenite and sorbitol as compared to cytoplasmic RNA) in U2OS cell line were used (GEO identifiers: GSE138988, GSE99304, GSE119977) ^44–46^. In detail, the following SRA (Sequence Reads Archives) data were employed (GEO identifier [SRA codes]): GSE138988[SRR10294571, SRR10294572, SRR10294573, SRR10294574, SRR10294575, SRR10294576, SRR10294577, SRR10294578, SRR10294579, SRR10294580, SRR10294581, SRR10294582]; GSE99304[SRR5605163, SRR5605164, SRR5605165, SRR5605166, SRR5605167, SRR5605168]; GSE119977[SRR7830171, RR7830172, SRR7830173, SRR7830174, SRR7830175, SRR7830176, SRR7830177, SRR7830178, SRR7830179, SRR7830180, SRR7830187, SRR7830188, SRR7830189, SRR7830193, SRR7830194, SRR7830195, SRR7830196, SRR7830197, SRR7830198].

All Fastq were downloaded and de-duplicated (with FastUniq v1.1) and trimmed (with TrimGalore 0.6.10, default parameters). Alignments were performed with STAR (v2.7); read counts were obtained with HTseq(v2.0.3). The reference genome was the telomere comprehensive T2T-CHM13v2.0 T2T CHM13 (Telomere-to-Telomere) assembly (downloaded from https://github.com/marbl/CHM13; with the UCSC GENCODEv35 CAT/Liftoff v2 annotation file). The gff file was manually edited to include putative TERRA regions, at telomeric extremes of each chromosome, with the following strategy: the initial/final 15000 bases (from start: 1 to 15000; from end, end – 15000 positions) were considered as “TERRA regions”, unless a known gene/annotation was partially overlapping with it: in this case, the “TERRA region” was considered from *(i)* the start of the telomere to the beginning of the known annotation (e.g. position 7569 of the T2T assembly for chromosome 1), or *(ii)* the end of the known annotation to the end of the chromosome (e.g. from position 134752359, to the end of chromosome 10). With these definition, TERRA regions were then termed p followed by chromosome number (p1, p2, …., pX), for proximal (at the chromosome start) telomeric extremes, or q followed by chromosome number (q1, q2, …., qX) for distal telomeric extremes. Statistical analyses were performed with in-house R scripts (R version 4.3.3, edgeR v 4.0.16).

### Proteomic interactome analysis

#### Preparation of nuclear and cytoplasmic extracts

U2OS cells were harvested and incubated in hypotonic buffer (10 mM Hepes, pH 7.9, 1.5 mM MgCl2, 10 mM KCl) and homogenized with a glass douncer. The total cell extract was spun for 15 mins at 1500g at 4°C to pellet the nuclei, while the supernatant was harvested for use as cytoplasmic fraction. To prepare the nuclear fraction, the pelleted nuclei were washed with PBS and extracted in hypertonic buffer (420 mM NaCl, 20 mM Hepes, pH 7.9, 20% glycerol, 2 mM MgCl2, 0.2 mM EDTA, 0.1% Igepal CA630, 0.5 mM DTT) for 2 hours at 4°C on a rotating wheel.

#### RNA oligos

5’ -biotinlyated RNA oligos were synthesized by Integrated DNA Technologies as indicated below.

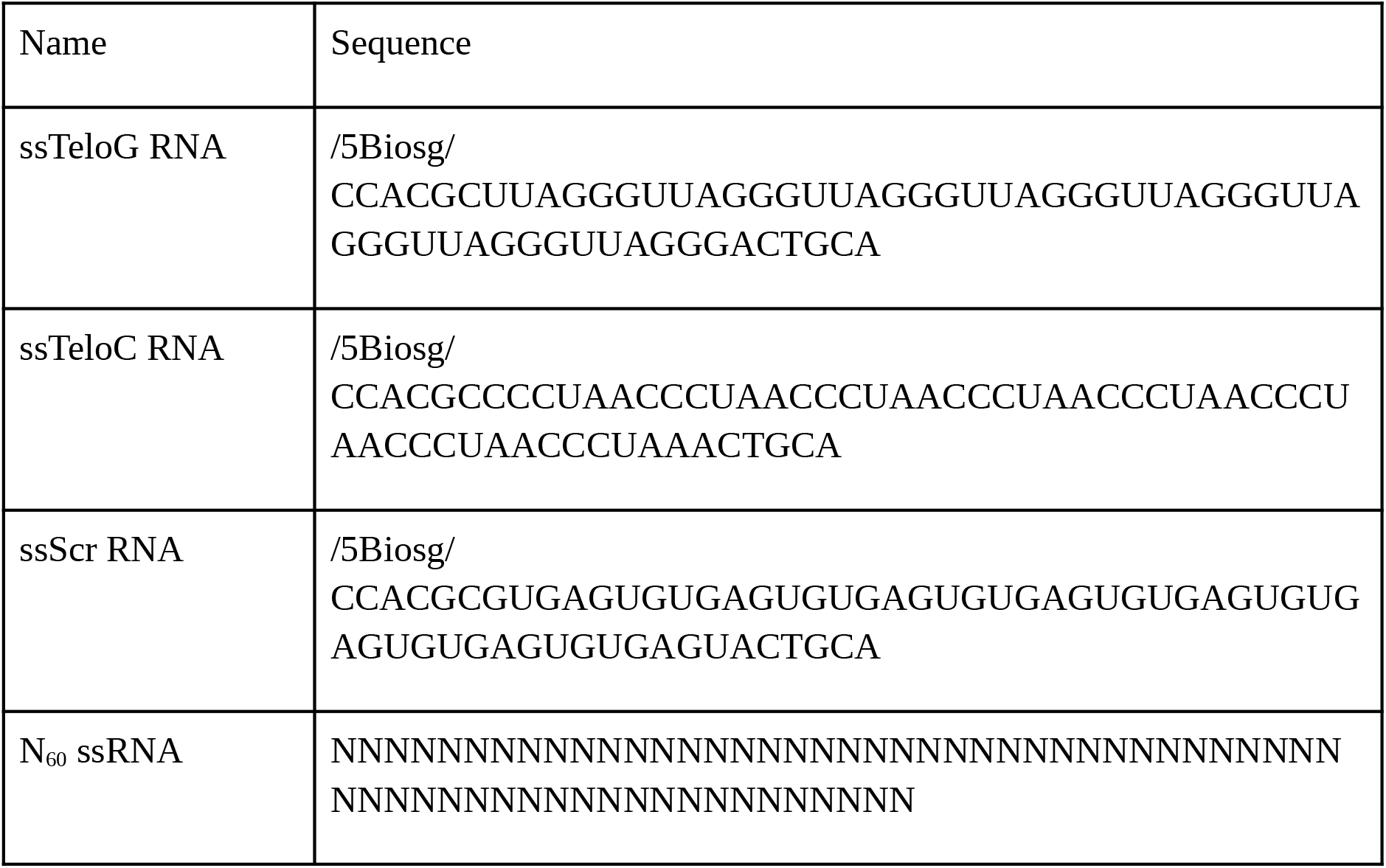

#### RNA Pulldown

25 µg of biotinylated RNA oligos were coupled to 25 µl streptavidin beads and incubated with 100 µg nuclear or cytoplasmic extracts in PBB buffer (150 mM NaCl, 50 mM Tris-HCl pH 7.5, 5 mM MgCl_2_, 0.5% Igepal CA-630) on a rotating wheel for 2 hours at 4°C. To quench non-sequence-specific interactions, 20 µg of N_60_ RNA oligo was added as a competitor for RNA binding. The beads were washed with PBB 3 times and boiled for 5 mins at 95°C in Laemmli buffer to elute the bound proteins.

#### Mass spectrometry

The pulldown elute was run on 12% Bis-Tris gel and cut into 2x2x1 mm pieces. The samples were reduced in 10 mM DTT for 1 hour at 56°C and then alkylated with 55 mM iodacetamide for 45 min in the dark. Samples were then digested with 2 µg trypsin in 50 mM ammonium bicarbonate buffer at 37°C overnight. The resulting peptides were subsequently desalted on StageTips and analyzed on an EASY-nLC 1200 Liquid Chromatograph coupled to a Q Exactive HF mass spectrometer via separation on a C18-reversed phase column (25 cm long, 75 μm inner diameter) packed in-house with ReproSil-Pur C18-AQ 1.9 μm resin (Dr Maisch) at 40°C. A 105-min gradient from 2 to 40% acetonitrile in 0.5% formic acid at 225 nl/min flow was used. Spray voltage was set to 2.2 kV. The Q Exactive HF was operated with a TOP20 MS/MS spectra acquisition method per MS full scan. MS scans were conducted with 60,000 resolution at a maximum injection time of 20 ms, and MS/MS scans with 15,000 resolution at a maximum injection time of 75 ms. The raw files were processed with MaxQuant version ^47^ 1.5.2.8 with preset standard settings for label-free quantitation with the MaxLFQ algorithm ^48^.

## Results

### LncRNA TERRA localizes to the cytoplasm of ALT cells

In order to investigate the presence of TERRA in the cytoplasm of human cells, we used single molecule inexpensive FISH (smiFISH), a sensitive variant of RNA-FISH ^41^. Using this technique, we successfully visualized cytoplasmic TERRA as clear foci in the osteosarcoma ALT cell line U2OS. Subsequently, we have found that this observation was reproducible for other two ALT cell lines: osteosarcoma cell line SA-OS2 and a lung-derived immortalized cell line WI-38 VA13 (Fig. 1A). While the cells of all three cell lines displayed high variability in the number of cytoplasmic TERRA foci, an RNase treatment decreased their numbers significantly (Fig. 1B). Conversely, we did not detect cytoplasmic TERRA in telomerase-positive, stomach adenocarcinoma cell line AGS (Supplementary Fig. 1), indicating that in telomerase-positive cancer cells, TERRA is predominantly nuclear or its levels are too low to enable detection of the cytoplasmic fraction.

**Fig. 1:**
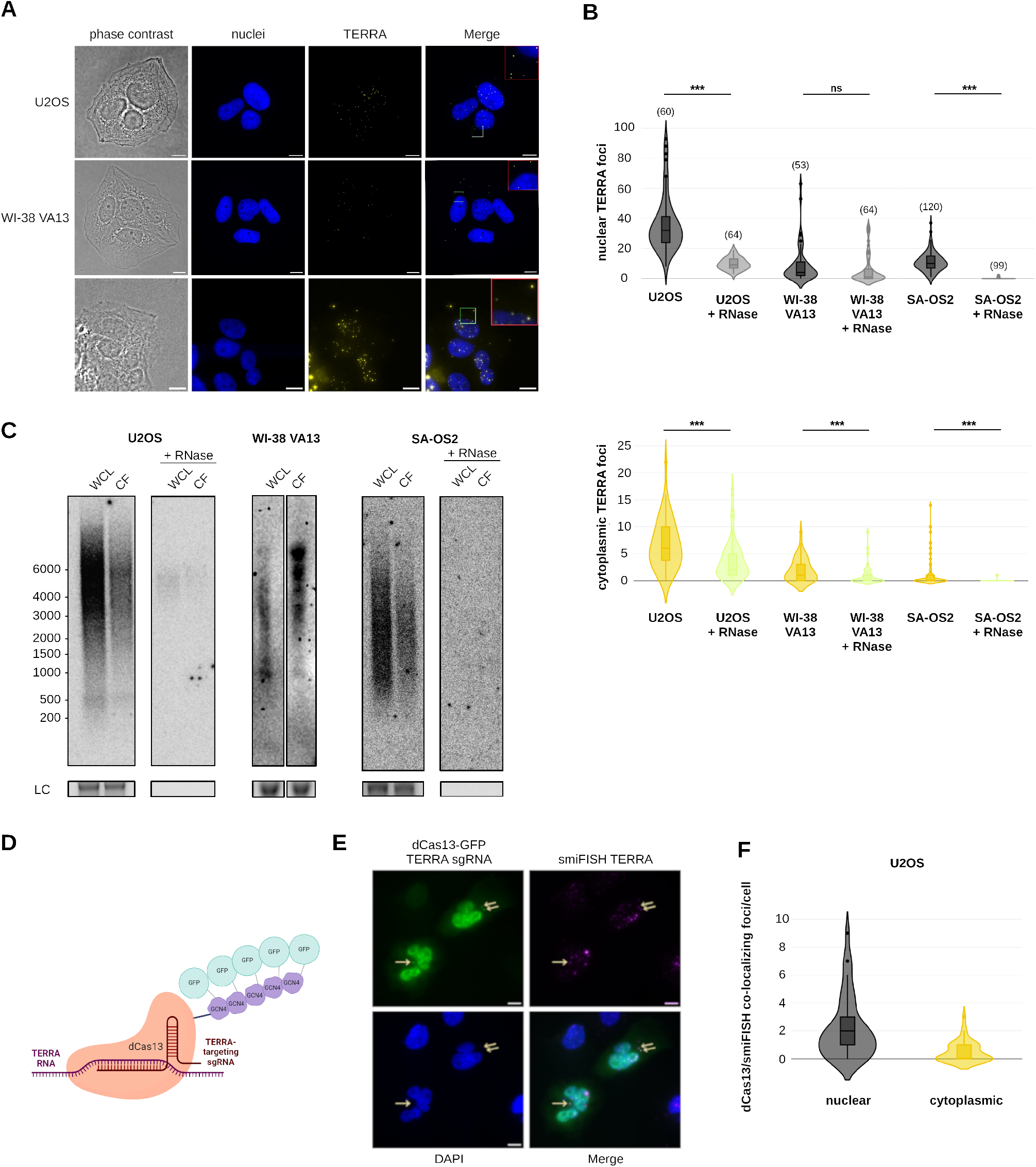
TERRA localizes to the cytoplasm in ALT cell lines. **A.** Representative pictures of U2OS, SA-OS2 ALT cancer cell lines, and WI38 VA13 immortalized ALT cell line displaying TERRA foci (yellow) both in the nucleus and in the cytoplasm. Nuclei were stained by DAPI (blue), cell borders were identified from phase contrast images. Scale bar: 10 µm. Insets (red square) are 3x enlarged selected regions (green square). **B.** Quantification of signal displayed in A, separately for nuclear and cytoplasmic foci. The impact of RNase A + RNase T1 (RNase) treatment is shown. The number of analyzed cells is displayed above each plot (the same cells were analyzed for both nuclear and cytoplasmic focus counts). ***: p<0.001, n.s.: not significant; unpaired two-tailed t-test was used to compare the samples. **C.** Northern blot analysis with ^32^P-(CCCUAA)_5_ probe demonstrates the presence of TERRA signal also in cytoplasmic fractions, with a similar distribution as can be seen in the whole cell lysate (WCL) sample. Same amount of RNA was loaded for the WCL and CF sample. 28S rRNA is shown as a loading control (LC). **D.** Scheme of the dCas13/sgTERRA complex that was used to detect TERRA. Five GFP molecules bind to the dCas13 through SunTag. Created with Biorender.com. **E.** An example of cytoplasmic foci that are positive for both dCas13-GFP/sgTERRA and TERRA smiFISH signal. CytoTERRA foci are highlighted by arrows. dCas13 fused with several nuclear localization sequence (NLS) was used for the experiments, to ensure low background in the cytoplasm, leading to high, diffuse background in the nucleus. Scale bar 10 µm. **F.** Quantification of nuclear and cytoplasmic dCas13-GFP/smiFISH double-positive foci in U2OS cells. 38 cells with at least one nuclear or cytoplasmic focus from 3 biological replicas were assessed for the quantification.

Previously, TERRA has been found in the extracellular exosomes in the form of a short, ∼200nt-long RNA mainly in human lymphoblastoid cell lines, especially during serum starvation ^35^. To further investigate whether the population of TERRA we observe in the cytoplasm (cytoTERRA) follows the same size distribution, we performed cell fractionation followed by a Northern blot with a probe complementary to the telomeric sequence of TERRA (Supplementary Fig. 2; Fig 1C). Interestingly, we found cytoTERRA ranging in size from 200 nt to more than 6 kb, similarly to nuclear TERRA ^11, 12^, suggesting that this population is more similar to the nuclear TERRA than the exosomal TERRA species. Additionally, we observed no change in the number of cytoTERRA foci per cell upon serum starvation (Supplementary Fig. 3).

To further confirm the presence of TERRA in the cytoplasm in U2OS cells, we employed a Cas-based visualization approach developed by Xu et al. ^38^. This method allows for a specific visualization of TERRA by catalytically-inactive RNA-binding Cas13 coupled with GFP through SunTag repeats (dCas13-GFP) and an sgRNA targeting telomeric sequence (sgTERRA) (Fig. 1D). In our experiments, this system exhibited a high specificity of binding, comparable with the original study—as more than 90% of dCas13-GFP TERRA foci colocalized with smiFISH TERRA signal in fixed cells ^38^(Supplementary Fig. 4A). While the sensitivity of this technique is lower than the smiFISH approach, as less TERRA foci can be visualizes using the dCas13-GFP system, we detected both nuclear and cytoplasmic TERRA foci that were positive for both dCas13-GFP/sgTERRA and smiFISH TERRA signal (Fig. 1E, F). Additionally, FACS sorting of GFP-positive cells yielded U2OS cells with higher levels of cytoTERRA as measured by dCas13-GFP/smiFISH co-localization events and corroborated the high variability of the number of cytoplasmic TERRA foci detected per cell (Supplementary Fig. 4C, D).

Overall, these results indicate the presence of cytoplasmic TERRA molecules in ALT cancer cells. CytoTERRA transcripts display similar length pattern as nuclear TERRA, suggesting that they represent full length transcripts, potentially actively exported from the nucleus.

### DNA damage at telomeres correlates with the levels of cytoTERRA

ALT cells are characterized by ongoing genome instability ^27^. Additionally, certain types of telomere dysfunction lead to an increase in total TERRA levels, including the loss of telomeric dsDNA-binding protein, TRF2, from telomeres ^21^. Based on these observations, we hypothesized that the intercellular variability in cytoplasmic TERRA levels may be explained by different amount of telomeric DNA damage accumulated in individual cells. To investigate this possibility, we assessed the presence of cytoTERRA and the amount of telomere DNA damage in unperturbed ALT cells by smiFISH coupled with immunofluorescence (IF), considering the co-localization between a telomeric protein RAP1 and a marker of DNA damage, γH2AX, as a telomere dysfunction-induced focus (TIF), a marker of telomeric DNA damage. Using this approach, we found a correlation between the number of TIFs with nuclear TERRA levels, but also with cytoplasmic TERRA levels in WI3-8 VA13 cell line (Supplementary Fig. 5), supporting the hypothesis that DNA damage at telomeres may be linked to the accumulation of cytoTERRA.

To further explore the connection between telomeric DNA damage and cytoTERRA, we depleted TRF2 protein from telomeres by overexpressing its dominant-negative allelle, TRF2ΔBΔM, in U2OS cells ^49^, which resulted in increased number of nuclear TERRA foci as expected due to the higher total TERRA levels reported upon TRF2 depletion ^21^. Furthermore, we observed that the TRF2ΔBΔM-expressing cells displayed an accumulation of the cytoplasmic TERRA foci, supporting the hypothesis that cytoTERRA is increased in the presence of telomeric DNA damage (Fig. 2A).

**Fig. 2:**
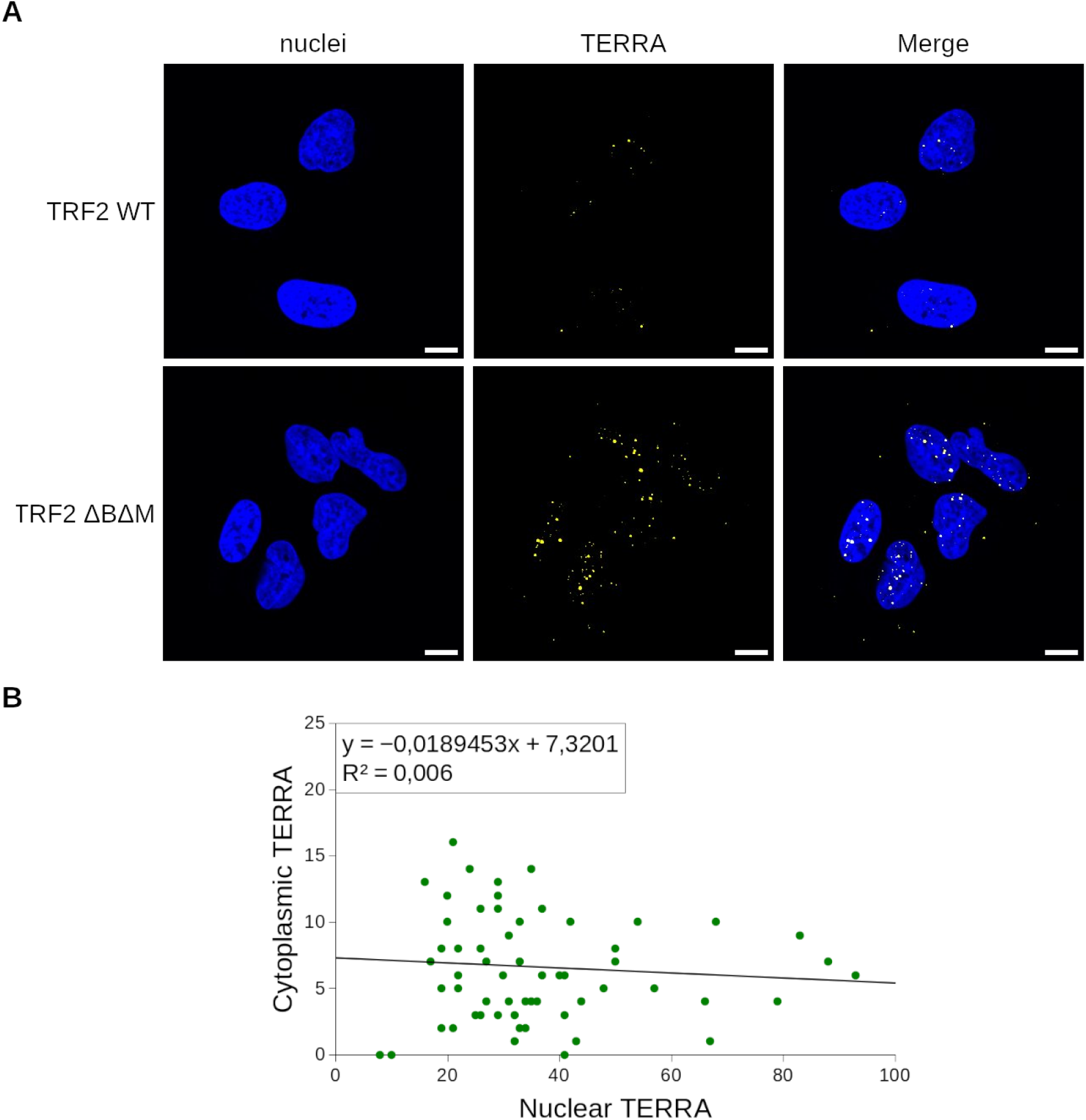
Overexpression of dominant negative form of TRF2 increases both nuclear TERRA and cytoTERRA foci numbers. **A.** Representative images of TERRA signal as detected by smiFISH performed inTRF2 WT and TRF2 ΔBΔM-transduced U2OS cells, respectively. Nuclei were stained by DAPI (blue). Scale bar: 10 µm. Images are representative of 2 biological replicas. **B.** Nuclear and cytoplasmic TERRA levels do not correlate in unperturbed U2OS cells. A linear regression analysis was performed on TERRA foci in untreated U2OS cells.

The absence of TRF2 at telomeres induces telomere deprotection through alteration of telomere structure and ATM signalling activation ^50, 51^. To assess whether different sources of telomeric damage would also lead to cytoTERRA accumulation, we depleted the telomeric ssDNA-binding protein POT1 in U2OS cells. POT1 depletion leads to ATR signalling activation and telomere instability driven by homology-directed repair, but unlike TRF2 depletion, it does not influence TERRA levels ^21, 51, 52^. Interestingly, the depletion of POT1 did not yield increased numbers of cytoTERRA foci (Supplementary Fig. 6), indicating a nuanced relationship between telomere homeostasis and export of TERRA to the cytoplasm.

Importantly, a quantitative analysis of individual unperturbed U2OS cells shows that the number of nuclear TERRA foci does not correlate with the number of cytoplasmic TERRA foci (Fig. 2B), supporting the hypothesis that elevated nuclear TERRA levels do not simply determine elevated cytoTERRA levels. These findings suggest that TERRA is not released to the cytoplasm upon its accumulation in the nucleus, but may be exported in an active fashion.

### TERRA interacts with nuclear export protein NXF1

To gain information on the potential function of cytoTERRA, its export and precise localization, we decided to perform a systematic incteractomics analysis similar to a previous study that characterized nuclear TERRA protein interactors ^53^. Here, a 60 nt-long, biotinylated TERRA RNA oligonucleotide was incubated with either nuclear or cytoplasmic cellular fractions, and the pulled-down proteins were identified by label-free quantitative mass spectrometry (MS) and compared to analogous pull-downs with a C-rich telomeric RNA or a scrambled control of the TERRA RNA. Among the identified candidates, we detected the members of the nuclear RNA export heterodimer NXF1/NXT1 interacting with TERRA oligonucleotide in both nuclear and cytoplasmic fraction (Fig. 3A). NXF1/NXT1 (also known as TAP/p15) heterodimer interacts with nuclear pore and drives the nuclear export of mRNAs ^54, 55^. NXF1 has been shown to be involved in the nuclear export of lncRNAs, especially transcripts with few exons, stable secondary structures and m6A modification ^4^, making TERRA a candidate for NXF1-mediated transport. To further investigate this possibility, we depleted NXF1 in U2OS cells via shRNA and assessed cytoTERRA levels in these cells compared to a scramble control (Fig. 3B–D). This experiment showed that the reduced levels of NXF1 lead to decreased detection of cytoTERRA by smiFISH as compared to scramble control, supporting the scenario in which TERRA is not passively transported to the cytoplasm upon the increase of nuclear TERRA levels, but its export to the cytoplasm is at least partially driven by the NXF1 export pathway.

**Fig. 3:**
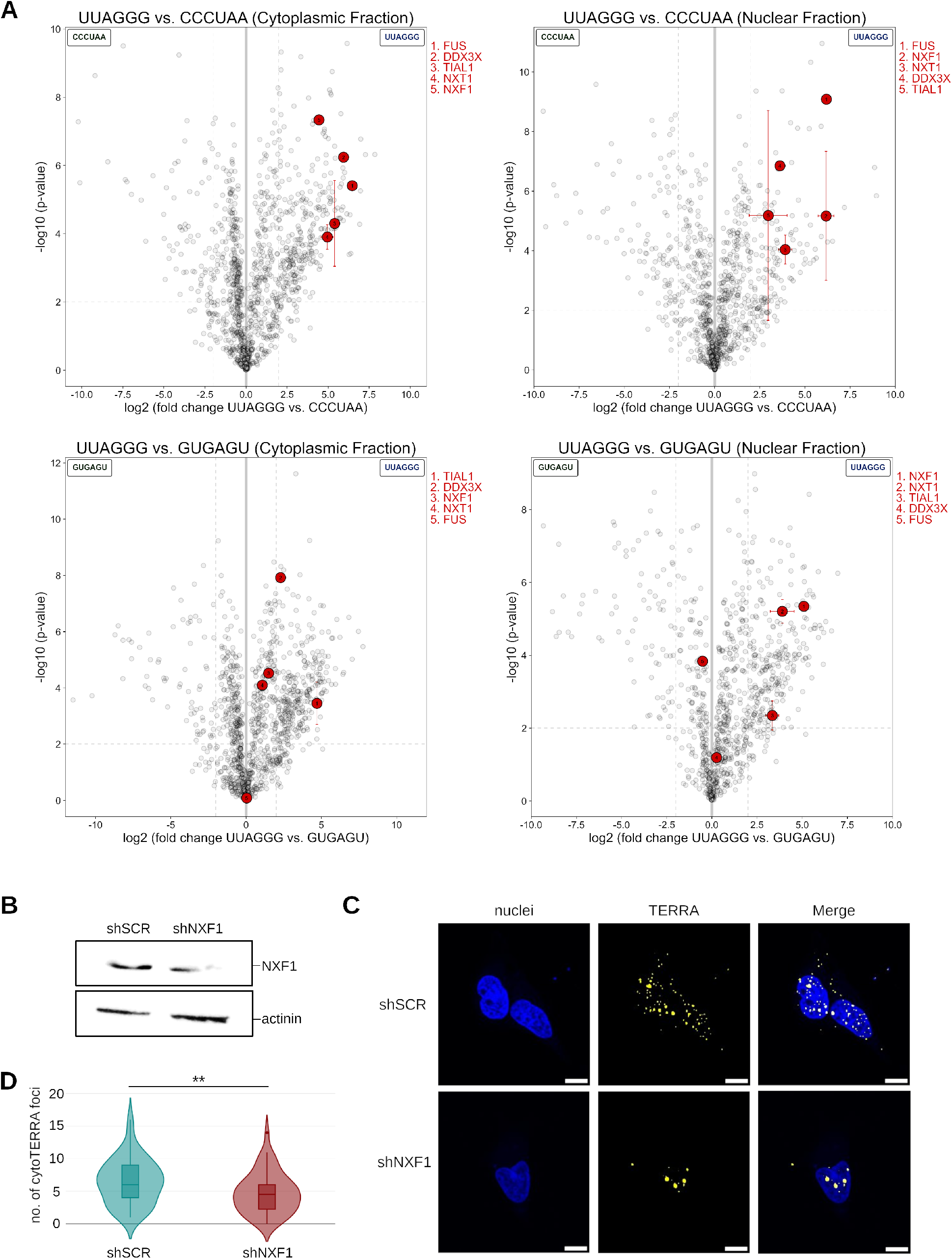
Loss of NXF1 leads to a decrease in cytoTERRA levels. **A.** Proteomics analysis identified a number of nuclear and cytoplasmic factors that interact with TERRA RNA. Volcano plots display proteins bound by G-rich telomeric sequence, as compared to a complementary C-rich or a scramble G-rich control, analyzed in cytoplasmic and nuclear fraction separately. Five proteins, including NXF1 and SG proteins FUS, DDX3X and TIAL1, are highlighted in the plots. **B.** Partial depletion of NXF1 by shRNA was assessed by Western blotting. Displayed is a representative image from 2 biological replicas. **C.** Representative image of shSCR-transfected vs. shNXF1-transfected U2OS cells. Scale bar: 10 µm. **D.** Quantification of cytoTERRA levels in shSCR and shNXF1-transfected U2OS cells. 51 and 53 cells, respectively, from 2 biological replicas were analyzed. **: p<0.01.

### CytoTERRA is increased upon stress and localizes to stress granules

Our experiments described above suggest that cytoTERRA is exported to the cytoplasm of ALT cells, possibly upon the accumulation of telomeric DNA damage. Next, we set out to gain further insight into the precise localization of cytoTERRA.

Our proteomics TERRA interactome revealed a number of proteins associating with TERRA described to localize to stress granules (SGs) (Fig. 3A). SGs are membrane-less organelles, forming in the cytoplasm of mammalian cells under different types of stress and serve to sequester the components of stalled translation machinery, such as translation initiation factors, ribosomal proteins, RNA and RNA-binding proteins, during translation inhibition and release them once the stress subsides ^8^.

To further investigate the connection with SGs, we induced stress in U2OS cells via two ways described previously to trigger the SG formation: sorbitol treatment that induces hyperosmotic stress and sodium arsenite, inducing oxidative stress. In sorbitol-treated cells, cytoTERRA was found to co-localize with FUS and TIAL1, as assessed by smiFISH/IF (Supplementary Fig. 7). Next, we used G3BP1 as a marker of SGs—it is a highly abundant SG component and was not identified as an interactor of TERRA in our proteomics screen. We observed a co-localization of a fraction of cytoTERRA foci with G3BP1 (Fig. 4A). The quantification of these colocalization events showed that upon sorbitol treatment, around ∼20% of cytoTERRA foci co-localize with SGs (Fig. 4B). Interestingly, the overall number of cytoTERRA foci is also increased in sorbitol compared to previous experiments in untreated cells, achieving levels comparable with nuclear TERRA foci (Fig. 4B).

**Fig. 4:**
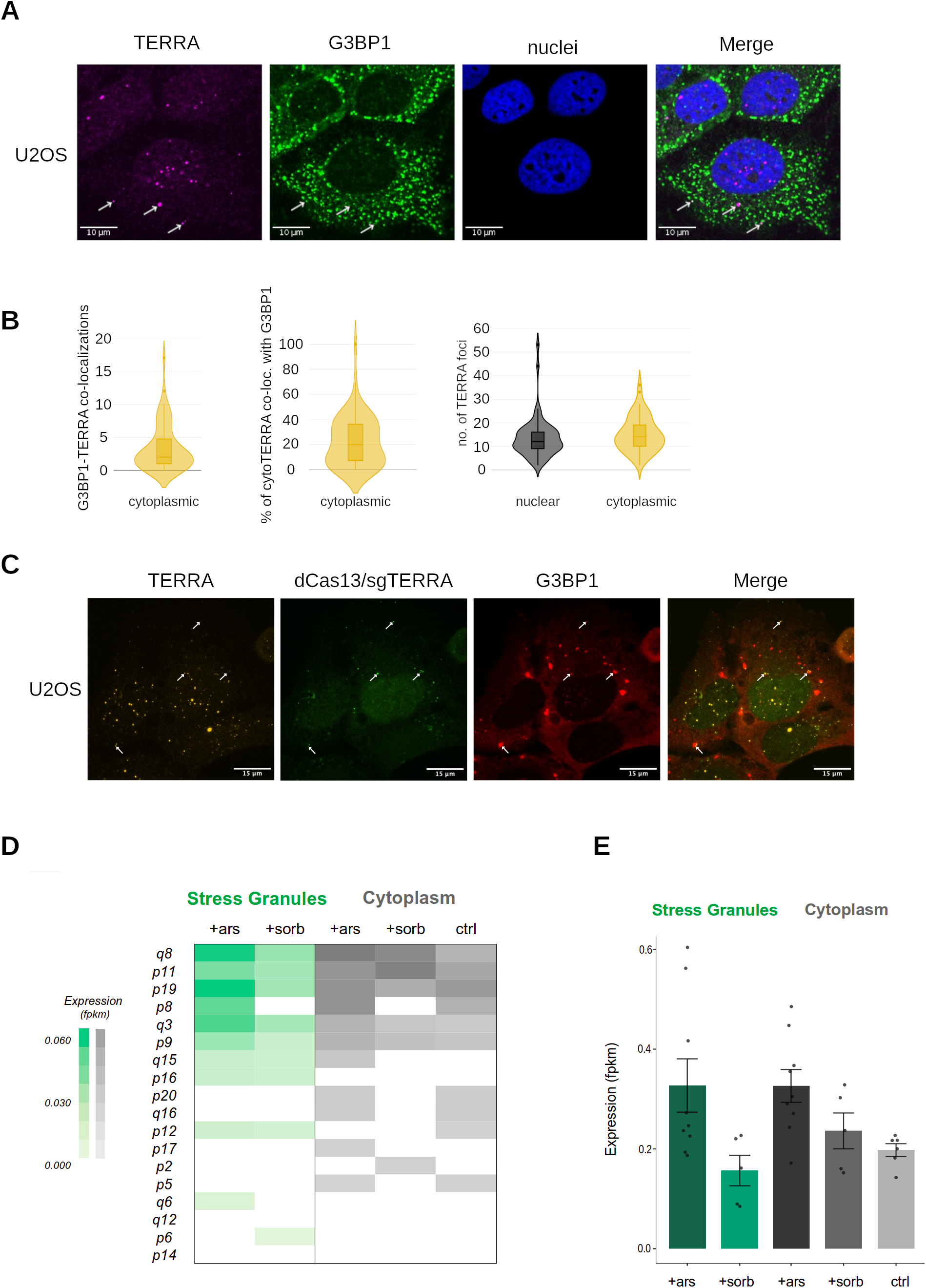
CytoTERRA localizes to stress granules. **A.** Representative image of co-localization events between cytoTERRA and G3BP1, a stress granule protein, under sorbitol treatment in U2OS cells, as assessed by smiFISH/IF experiments. Scale bar: 10 µm. Considering the high number of G3BP1 foci, the “shuffle” function in ImageJ DiAna plugin was used to corroborate that the colocalization detected is non-random. **B.** Quantification of the co-localization between TERRA and G3BP1 during sorbitol treatment, showing the absolute number of cytoTERRA-G3BP1 co-localization events and relative percentage of cytoTERRA foci co-localizing with G3BP1 per cell and the absolute numbers of foci in nucleus and cytoplasm. 58 cells were assessed for the analysis. **C.** An example of co-localization between TERRA and G3BP1 in U2OS cells in arsenite stress conditions as assessed by dCas13-GFP/smiFISH TERRA visualization. Co-localization events highlighted by arrows. Scale bar: 15 µm. **D.** The abundance of TERRA from individual subtelomeres detected in RNA-seq data reported for cytoplasm and stress granules of U2OS cells unstressed (ctrl) and treated with sorbitol (sorb) or arsenite (ars). Values reported refer to the average abundance across replicates (at least 3). Only TERRA elements with detectable expression levels in at least two thirds of the replicates are reported. **E.** Relative abundance of TERRA reads (fpkm) in the RNA-seq data in all conditions analyzed. Plotted is mean +- s.d., single replicates are displayed as points.

Using a smiFISH approach to observe cytoTERRA upon sodium arsenite treatment (oxidative stress), we did not detect co-localization with SGs or elevated cytoTERRA numbers in stressed cells compared to the control (data not shown). However, using the dCas13-sgTERRA/smiFISH combined cytoTERRA visualization approach upon sodium arsenite treatment, we observed that a proportion of cytoTERRA co-localizes with SGs, similarly to sorbitol treatment (Fig. 4C). This observation may suggest that the additional stress of dCas13-GFP/sgTERRA transfection combined with sodium arsenite-induced oxidative stress cumulatively result in increased cytoTERRA levels.

To use an orthogonal approach to assess the presence of TERRA at SGs, we analyzed publicly available SG transcriptome data in U2OS cells ^44–46^. We re-mapped the RNA-seq reads derived from isolated SGs under sorbitol and sodium arsenite stress and compared these results with the cytoplasmic transcriptome from stress and unstressed controls. In both sorbitol and sodium arsenite stress, our analysis identified TERRA reads in both the cytoplasmic and the SG transcriptome, as well as in the unstressed cytoplasmic control (Fig. 4D–E). Focusing on TERRA derived from individual chromosome ends, we found that there is no enrichment for a specific TERRA species at SGs compared to the corresponding cytoplasmic transcriptome from stressed conditions (Fig. 4D). The same subset of TERRA species—mainly telomere q8, p11, p19, p8, q3 and p9-derived TERRA—is overrepresented under stress in both cytoplasmic transcriptome and at SGs, although these are not necessarily the TERRA species most abundantly represented in the cytoplasm in steady-state conditions. These data suggest that TERRA is recruited to the SGs based on the presence of its telomeric sequence that is identical for all TERRA species, not the subtelomere-specific 5’ sequence. Quantifying the relative representation of TERRA in all surveyed transcriptomes, we confirmed our observation that overall cytoTERRA levels increase upon sorbitol, and an increase was also detected in sodium arsenite stress (Fig. 4E). Interestingly, the relative abundance of TERRA at SGs is higher in sodium arsenite stress conditions than upon sorbitol treatment (Fig. 4E). This may reflect absolute levels of TERRA, but may also be a result of other RNA species being highly abundant in the sorbitol, but not sodium arsenite SG transcriptome.

In conclusion, the combined results from the proteomics screen, microscopy co-localization analysis and the analysis of SG transcriptome suggest that during osmotic and oxidative stress, cytoTERRA is a component of stress granules in ALT cells and is recruited to SGs independently of its telomere of origin.

## Discussion

Several recent reports found TERRA outside the nucleus or hinted at its possible cytoplasmic localization—the demonstration of TERRA in extracellular vesicles ^34, 35^, the discovery of its involvement in the senescence-triggered inflammation signalling at mitochondria ^36^ or the investigation of its translation potential ^37, 56^.

In this study, we show with several independent techniques that TERRA is present in the cytoplasm of ALT cells and we term this population cytoTERRA. Employing sensitive microscopy techniques, cellular fractionation, proteomics analysis and re-mapping existing SG RNA-seq datasets, we show that the numbers of cytoTERRA foci increase upon telomeric DNA damage, osmotic and oxidative stress. Furthermore, we provide evidence that the nuclear export of TERRA is partially mediated by NXF1 nuclear export pathway and that a fraction of cytoTERRA is localized at stress granules.

It is interesting to speculate about the role of TERRA in stress granules. One possibility is that it may contribute to their formation—indeed, G-quadruplex forming RNAs (RG4s) were previously found to stimulate the phase separation of these bodies ^57, 58^. Additionally, not only TERRA readily forms rG4 in vitro and in cellulo ^19, 59, 60^, but a recent work has also demonstrated that stress conditions (such as starvation and sodium arsenite exposure) stabilize otherwise transient rG4 structures in the cytoplasm (^61^, Fig. 5A).

**Fig. 5:**
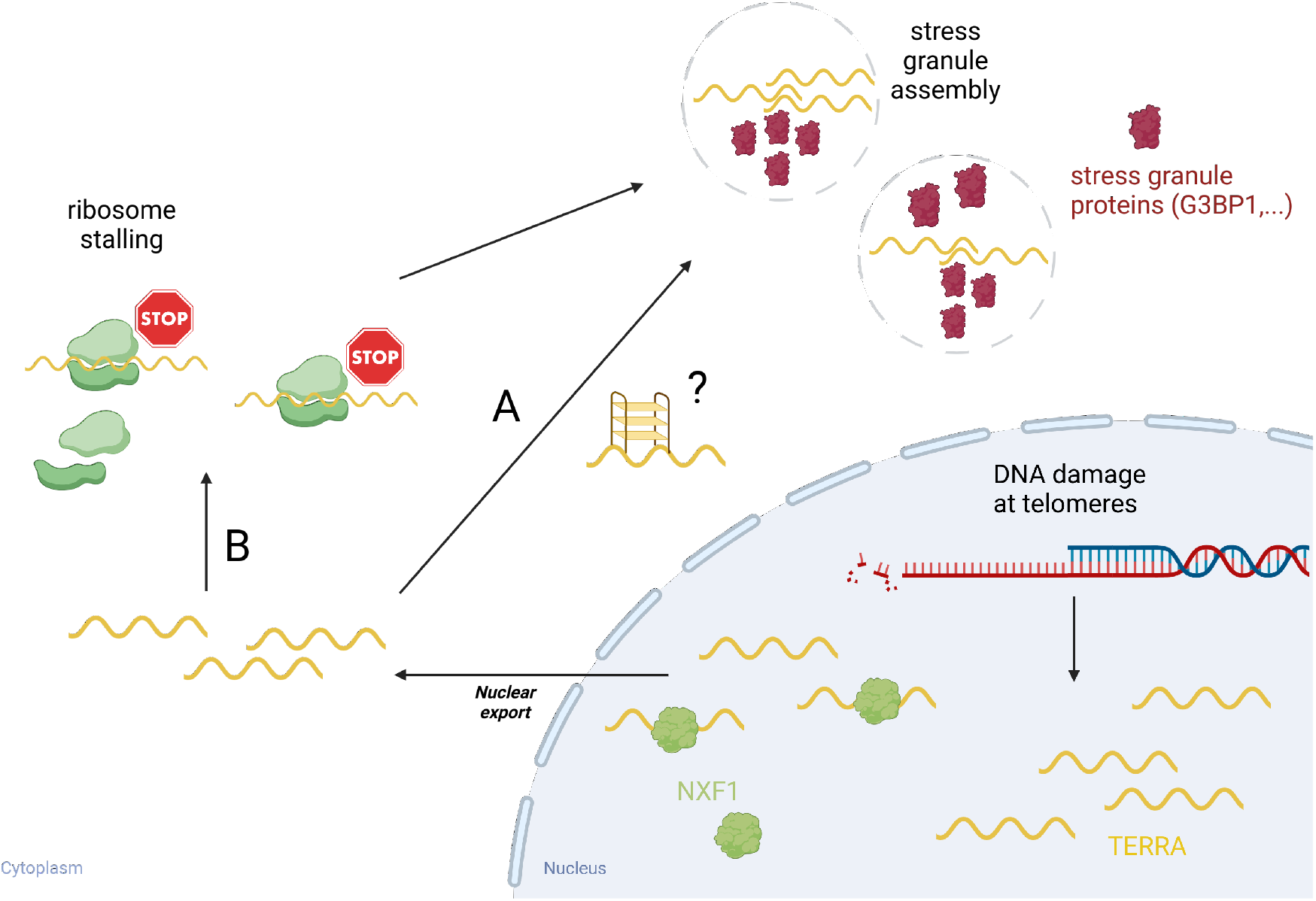
Proposed model of cytoplasmic TERRA roles. We hypothesize that upon its export from nucleus mediated by NXF1, cytoTERRA may associate with stress granules via two routes: A) directly, hypothetically contributing to the stress granule assembly—a process that may be favored by RNA G-quadruplex formation or B) via productive or unproductive binding to ribosomes that may be followed by translational stalling and sequestration of translation machinery factors and TERRA in the stress granules (see Discussion for details). Created with Biorender.com.

As SGs are typically formed upon translational stress, it is worth considering whether TERRA may first bind to ribosomes and only subsequently localize to SGs, similarly to other translated RNAs. Interestingly, our GO term analysis of the cytoplasmic interaction partners of TERRA showed that it has a capacity to interact with a number of translation initiation factors (Supplementary Fig. 8). Bringing this observation together with demonstration that TERRA may be translated ^37^, it suggests a possibility that the association of TERRA with SGs is preceded by productive or unproductive interaction with translation machinery (Fig. 5B).

In summary, the presence of TERRA at SGs opens a possibility that TERRA is involved in the general cellular response to stress, including telomeric DNA damage accumulation, that involves translation inhibition. It would be intriguing to elucidate whether cytoTERRA plays an active role in stress signalling and stress response.

Lastly, it is important to acknowledge that more insight will be necessary to fully understand the export of TERRA from nucleus to the cytoplasm and its connection to DNA damage at telomeres. While our experiments suggest the involvement of NXF1, it is possible that a fraction of TERRA remains cytoplasmic upon nuclear membrane disintegration during cellular division, requiring no active export from the nucleus. Additionally, it would be interesting to describe the regulation of the nuclear export of TERRA, to better understand whether it is simply increased levels of nuclear TERRA that drive the increase in its export, or whether other factors are at play. Finally, as ALT cells exhibit increased levels of global TERRA ^26–30^, it would be important to dissect whether these increased levels of TERRA simply enable for efficient visualization of cytoTERRA in ALT cells or whether the presence of increased levels of TERRA enable ALT cells to deploy it for its cytoplasmic roles upon telomeric DNA damage accumulation.

## Conclusion

In this work, we directly demonstrate the presence of TERRA in the cytoplasm of human telomerase-negative, ALT cells. We also show its levels are increased upon telomeric DNA damage accumulation, oxidative and osmotic stress, and that a fraction of cytoplasmic TERRA associates with stress granules. This study opens a number of questions about the function and relevance of cytoTERRA for nucleo-cytoplasmic communication during stress. We believe that this work contributes to a better understanding of the extranuclear functions of this multifaceted lncRNA.

## Acknowledgements

We would like to acknowledge Madan Mohan Udaya Kumar and Dennis Kappei (National University of Singapore) for providing expertise and technical support for the MS analysis of TERRA interactome. and critical reading of the manuscript. We would like to thank past and present members of Cusanelli laboratory for fruitful discussions and the Advanced Imaging Facility at CIBIO (Giorgina Scarduelli and Michela Roccuzzo) for invaluable technical assistance. We are grateful to Huaiying Zhang and Meng Xu (Carnegie Mellon University) and Andrea Lunardi and Alessandro Alaimo (University of Trento) for reagent sharing. This work received funding from the European Union’s Horizon 2020 research and innovation programme under the Marie Sklodowska-Curie Grant agreement No. 101032702 and Fondazione Veronesi (KJ).

## Notes

### Competing Interest Statement

The authors have declared no competing interest.

