## Supplementary Information for "Telomeric lncRNA TERRA localizes to stress granules in human ALT cells"

**Supplementary Figures**

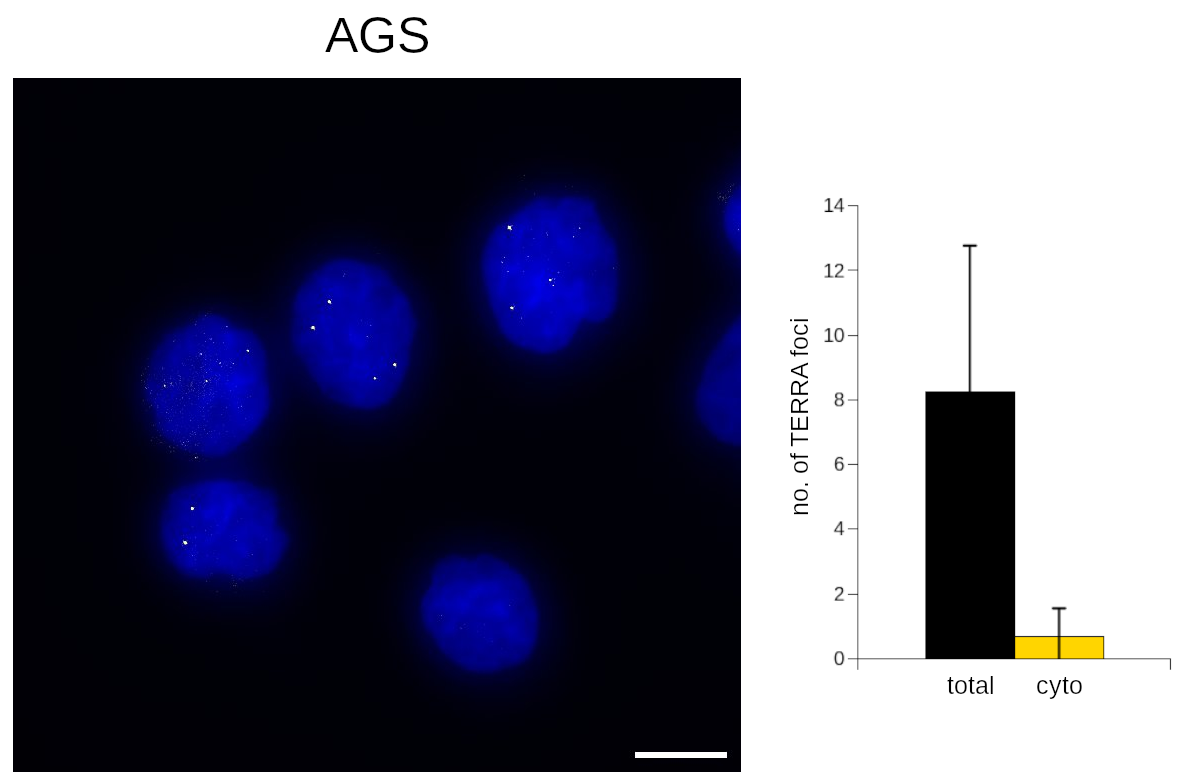

**Supplementary Figure 1: TERRA smiFISH in AGS.** Left: a representative image, DAPI: blue, TERRA smiFISH: yellow foci. Right: Quantification of total and cytoplasmic foci numbers. 28 cells were analyzed. Scale bar: 10 µm.

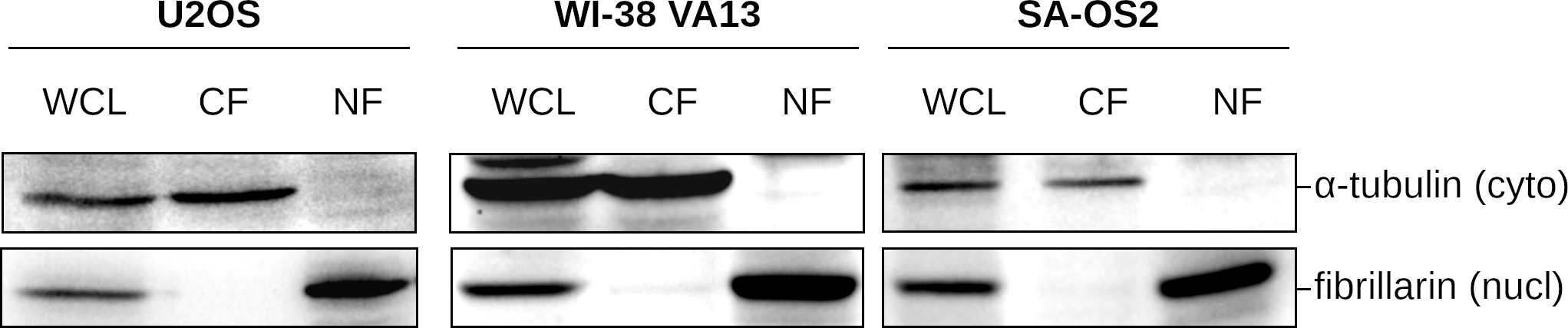

**Supplementary Figure 2:** **Western blot demonstrating the efficiency of fractionation for the three analyzed cell lines.** ɑ-tubulin was used as a cytoplasmic marker, fibrillarin as a nuclear marker. Whole cell lysate (WCL), cytoplasmic (CF) and nuclear (NF) fractions were analyzed.

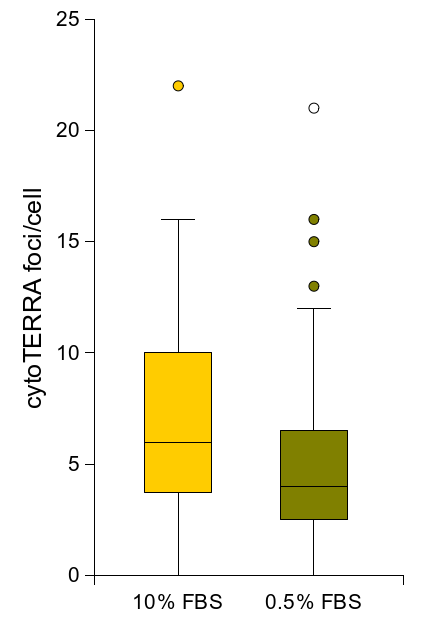

**Supplementary Figure 3: Serum starvation does not drive increase in cytoTERRA levels in U2OS cells.** Serum starvation was performed as described in (Wang et al. 2015). 60 and 63 cells, respectively, were quantified for the analysis.

**Su**
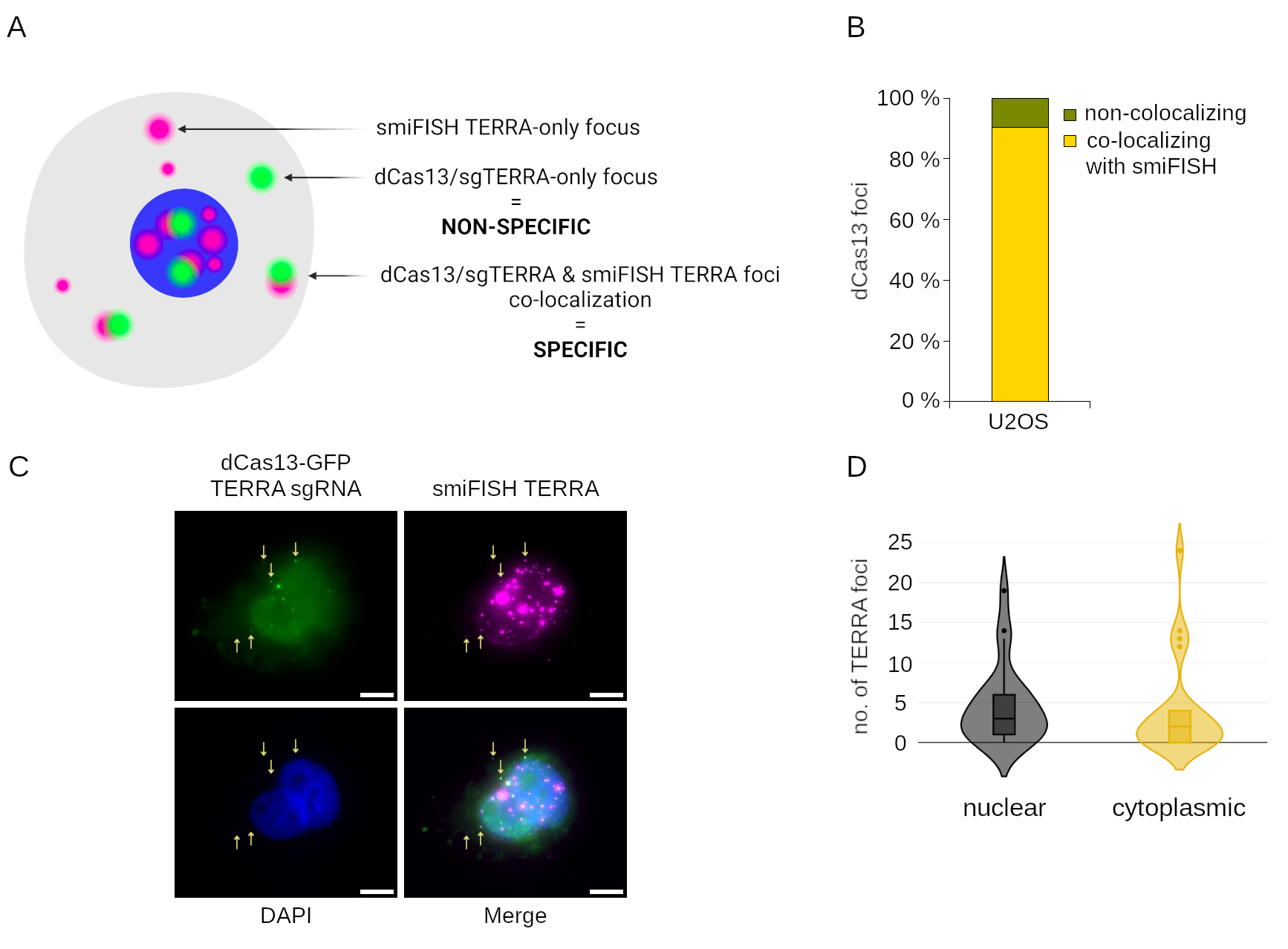
**pplementary Figure 4: dCas13 specificity and dCas13-GFP/sgTERRA signal in FACS-sorted cells. A**. Scheme of the dCas13 labeling specificity assessment. SmiFISH TERRA has a higher sensitivity than dCas13/sgTERRA, leading to more smiFISH-only foci (purple only). dCas13-GFP/sgTERRA-only foci (green only) were considered to be non-specific signal, possibly due to dCas13 aggregation. The co-localization of smiFISH TERRA and dCas13-GFP/sgTERRA foci (green & purple) was considered a specific visualization event. Created with Biorender.com. **B**. Quantification of specificity of dCas13-GFP/sgTERRA, visualized as a bar plot of the percentage of dCas13-GFP foci non-co-localizing and co-localizing with smiFISH foci out of all dCas13-GFP foci. 19 cells from two biological replicas were used for the analysis. **C.** Example of a dCas13-GFP/sgTERRA-expressing U2OS cell sorted by FACS as GFP+ and subjected to smiFISH TERRA (using ATTO-555 conjugated probe). Co-localizing dCas13-GFP/sgTERRA + smiFISH TERRA foci in the cytoplasm are highlighted by arrows. **D**. Quantification of specific (dCas13-GFP/sgTERRA & smiFISH TERRA) TERRA foci in FACS-sorted GFP+ cells; 18 cells were analyzed.

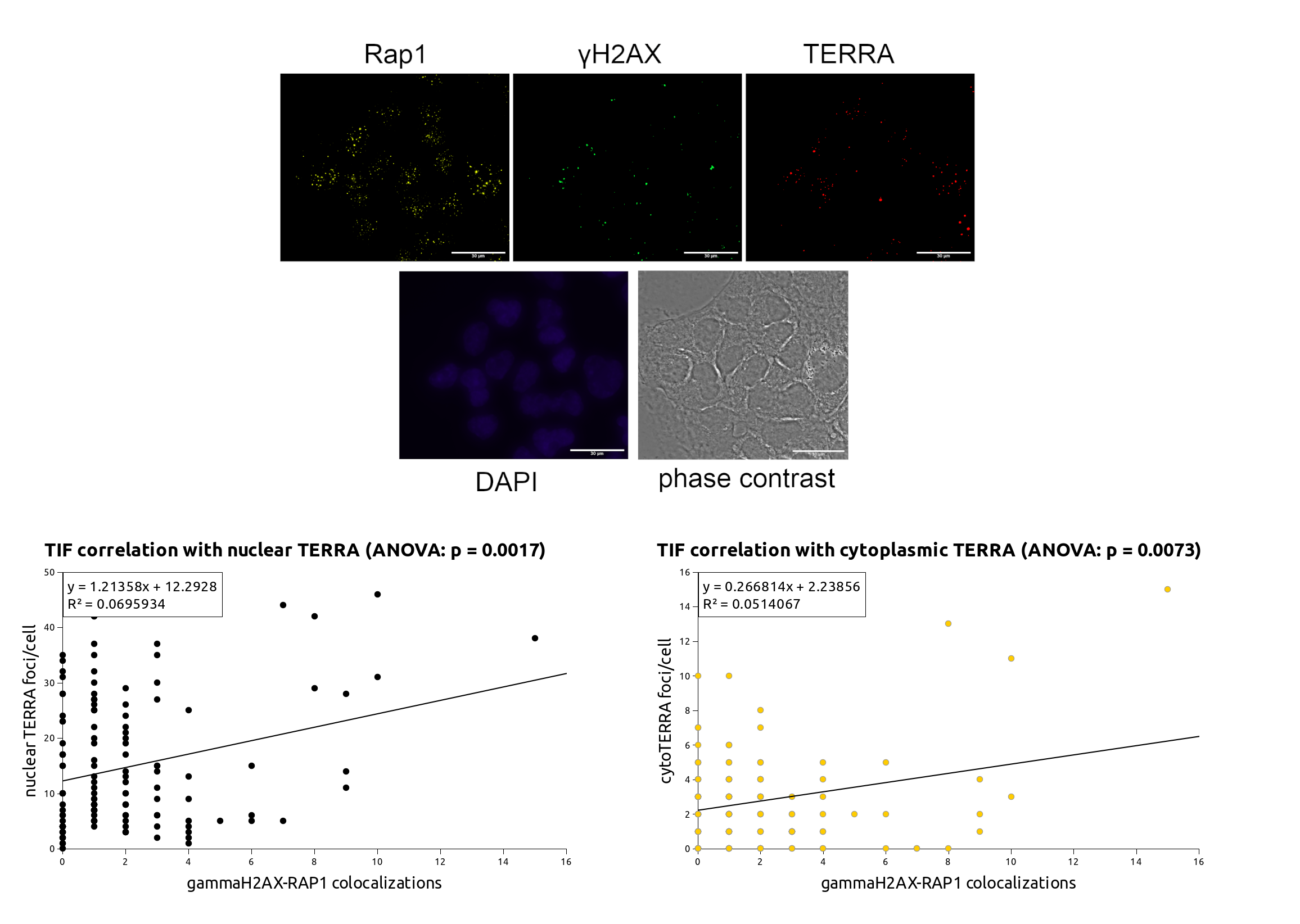
**Supplementary Figure 5: DNA damage at telomeres correlates with both nuclear and cytoplasmic TERRA.** Top: Representative image of WI-38 VA13 cells. Bottom: The correlation between the number of TIFs and TERRA foci was analyzed separately for nuclear and cytoplasmic TERRA foci. Multiple linear regression analysis was performed; 139 cells were analyzed.

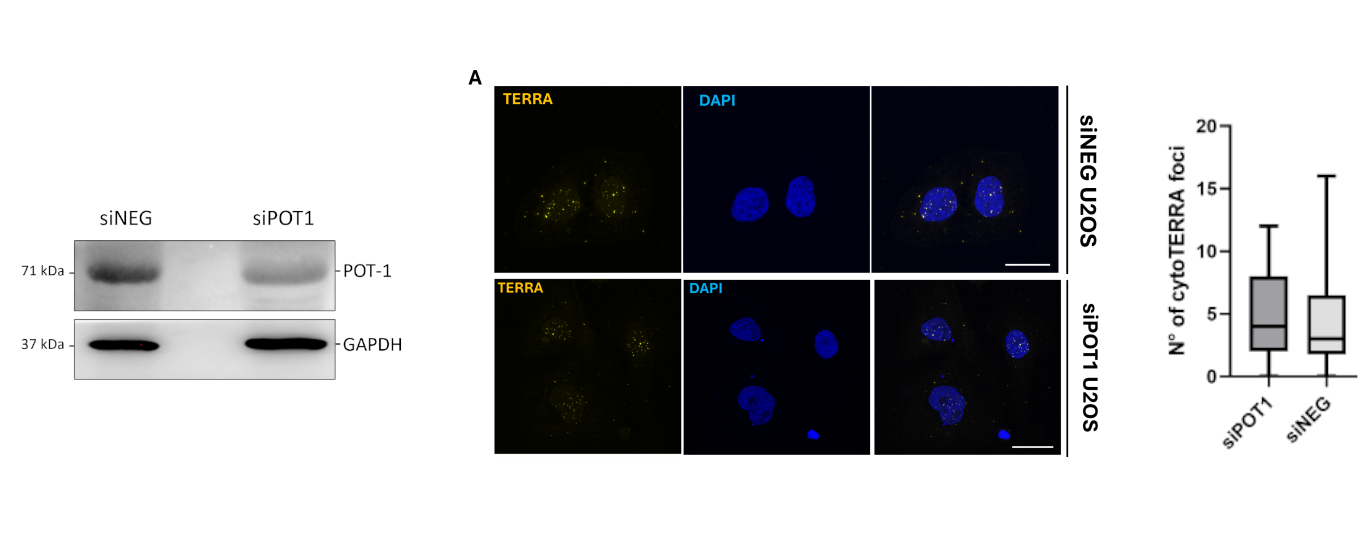

**Supplementary Figure 6: CytoTERRA levels are not impacted by POT1 depletion.** Left: Western blot demonstrates decreased levels of POT1 upon transfection of U2OS cells with POT1 targeting siRNA (siPOT1). Middle: Representative image of siNEG and siPOT1-treated cells. Right: Quantification of cytoTERRA levels.

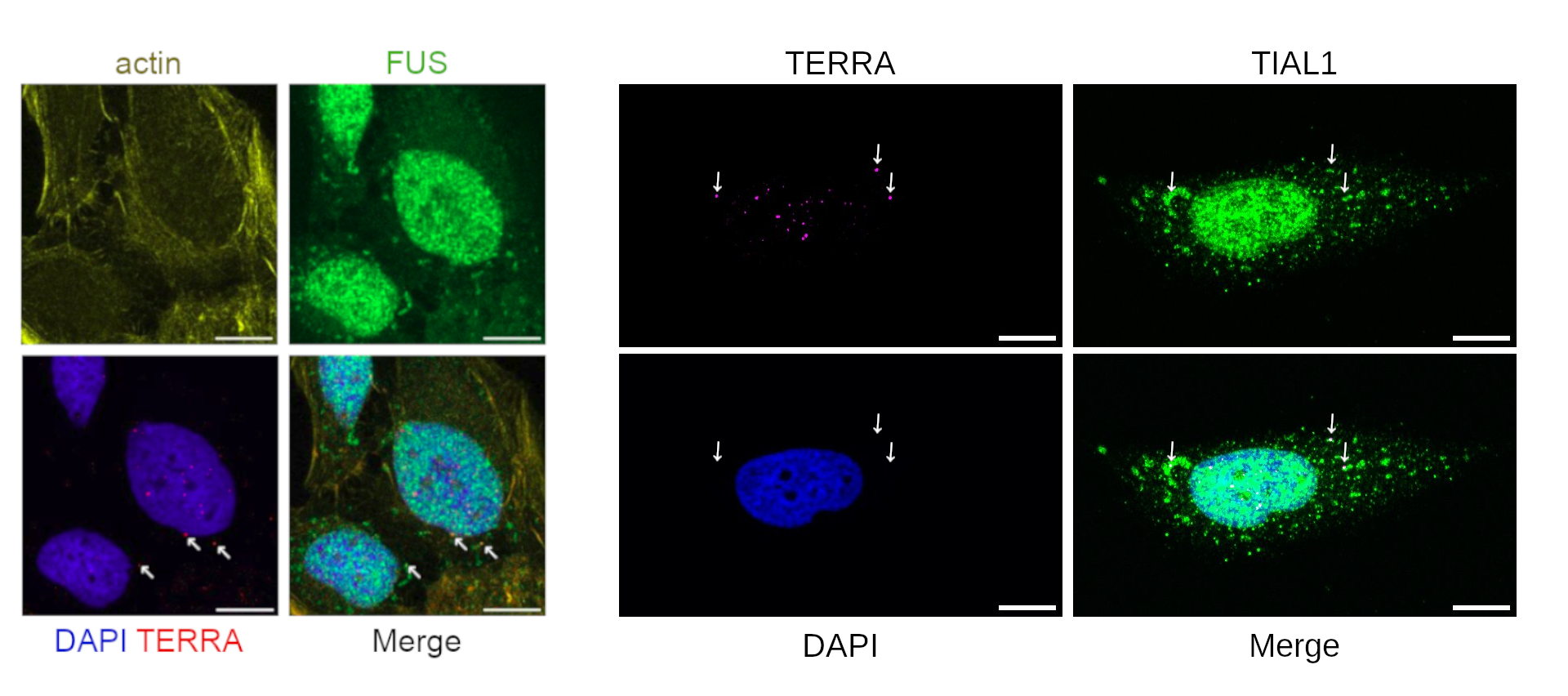

**Supplementary Figure 7: TERRA co-localizes with stress granule proteins FUS and TIAL1 in U2OS cells under sorbitol stress.** TERRA was visualized by smiFISH, with ATTO-555-conjugated probe; FUS and TIAL1 were visualized by IF, with Alexa488-conjugated secondary antibody. Representative images of are shown.

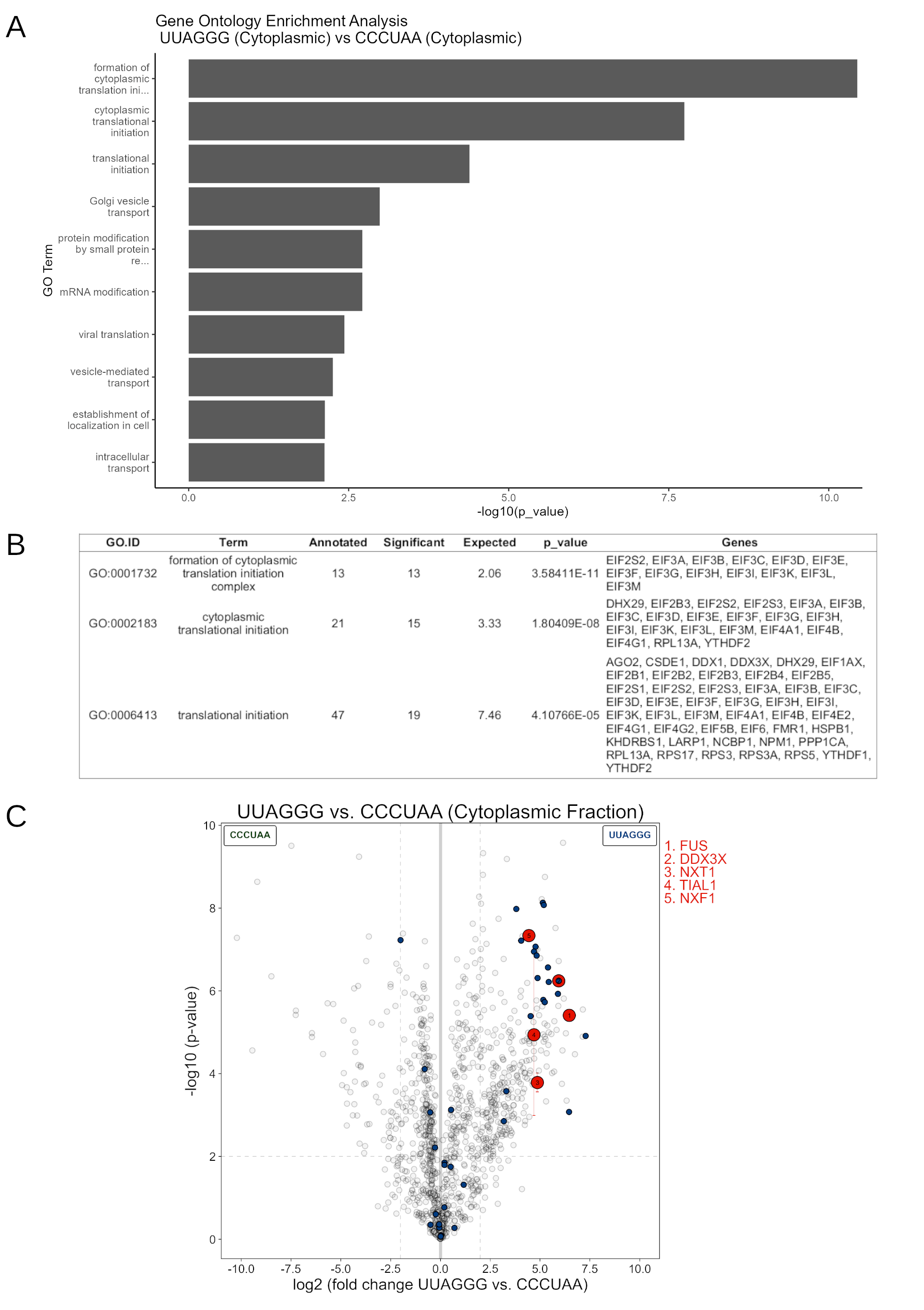

**Supplementary Figure 8: Protein interaction partners of TERRA include translation pre-initiation and initiation factors. A.**A GO (Biological process) enrichment analysis of cytoplasmic proteins interacting with TERRA-mimicking oligonucleotide when compared with the C-rich control is shown, demonstrating that the GO term "translation initiation" (GO:0006413) and two of its child terms were significantly enriched in the TeloG (cytoplasmic vs TeloC (cytoplasmic) comparison. **B.** Description table for the GO term enrichment analysis; the list of genes belonging in the translation initiation-related categories is shown. **C.** The volcano plot shown in Fig. 3A featuring the translation initiation-related factors highlighted in blue.
